# Considerations for reporting variants in novel candidate genes identified during clinical genomic testing

**DOI:** 10.1101/2024.02.05.579012

**Authors:** Jessica X. Chong, Seth I. Berger, Samantha Baxter, Erica Smith, Changrui Xiao, Daniel G. Calame, Megan H. Hawley, E. Andres Rivera-Munoz, Stephanie DiTroia, Genomics Research to Elucidate the Genetics of Rare Diseases (GREGoR) Consortium, Michael J. Bamshad, Heidi L. Rehm

**Author notes:** correspondence should be addressed to: Jessica Chong, PhD Department of Pediatrics University of Washington School of Medicine Box 357371 1959 NE Pacific Street, HSB I607I Seattle, WA 98195.

## Abstract

Since the first novel gene discovery for a Mendelian condition was made via exome sequencing (ES), the rapid increase in the number of genes known to underlie Mendelian conditions coupled with the adoption of exome (and more recently, genome) sequencing by diagnostic testing labs has changed the landscape of genomic testing for rare disease. Specifically, many individuals suspected to have a Mendelian condition are now routinely offered clinical ES. This commonly results in a precise genetic diagnosis but frequently overlooks the identification of novel candidate genes. Such candidates are also less likely to be identified in the absence of large-scale gene discovery research programs. Accordingly, clinical laboratories have both the opportunity, and some might argue a responsibility, to contribute to novel gene discovery which should in turn increase the diagnostic yield for many conditions. However, clinical diagnostic laboratories must necessarily balance priorities for throughput, turnaround time, cost efficiency, clinician preferences, and regulatory constraints, and often do not have the infrastructure or resources to effectively participate in either clinical translational or basic genome science research efforts. For these and other reasons, many laboratories have historically refrained from broadly sharing potentially pathogenic variants in novel genes via networks like Matchmaker Exchange, much less reporting such results to ordering providers. Efforts to report such results are further complicated by a lack of guidelines for clinical reporting and interpretation of variants in novel candidate genes. Nevertheless, there are myriad benefits for many stakeholders, including patients/families, clinicians, researchers, if clinical laboratories systematically and routinely identify, share, and report novel candidate genes. To facilitate this change in practice, we developed criteria for triaging, sharing, and reporting novel candidate genes that are most likely to be promptly validated as underlying a Mendelian condition and translated to use in clinical settings.

## Introduction

Over the past decade, phenotype-driven exome and genome sequencing (ES, GS), has demonstrated utility as a first-line test for clinical diagnostic testing in individuals with suspected Mendelian conditions.^1–5^ Nevertheless, a large fraction (∼30%-80%, depending on prior testing, categorical phenotype, analytical approach, and other factors) of persons tested by ES/GS remain without a precise genetic diagnosis,^3,6–11^ and a substantial proportion of these individuals may harbor variants in novel candidate genes that explain their clinical findings. While increasingly widespread application of newer technologies such as short-read genome sequencing,^12–14^ long-read sequencing,^15,16^ RNA-seq,^17,18^ and methylation analysis^19,20^ and accompanying analytical approaches/tools^21,22^ are facilitating new gene discoveries, most novel gene discoveries continue to be accessible with ES/GS.^23^ Furthermore, re-analysis of existing ES/GS data moderately improves diagnostic yield, with novel gene discoveries—i.e. newly-reported gene-disease relationships, of which an estimated ∼100-200 are published per year^24–26^— accounting for the majority of new precise genetic diagnoses.^8,23,27–30^ These findings highlight the value of incorporating recently described gene discoveries in the clinical analysis of ES/GS data, and suggest that even greater benefits may be leveraged by systematically sharing knowledge about candidate gene discoveries.

In contrast to single gene and multi-gene panel testing, ES/GS agnostically detects variants throughout the genome, including variants in genes that have been found to underlie a Mendelian condition (i.e., known genes) and genes that have not been implicated but for which varying sources of evidence may support a role in a Mendelian condition (i.e. novel candidate genes). As a result, if an individual’s clinical findings remain partially or completely unexplained after assessment of variants in known genes, assessing variants in novel candidate genes could result in a molecular diagnosis.^8,27–30^ Moreover, the increased cost (e.g., labor, time) to assess variants in novel candidate genes can be modest when implemented using computable criteria and added to an existing computational pipeline, suggesting a substantially increased return on the investment of generating the ES/GS data.

While extensive guidelines exist for assessing and reporting the pathogenicity of variants in known genes, including variants for which the clinical effect includes degrees of uncertainty,^31–37^ guidance is lacking for clinical interpretation and reporting of variants in novel candidate genes. We describe criteria to select novel candidate genes supported by sufficient evidence as to be of high clinical value, yet small enough in number so that routine inclusion in clinical ES/GS reporting at this time is manageable. Further, we provide guidance for reporting and sharing variants in novel candidate genes.

## Methods

### Drafting recommendations

We formed a working group within the GREGoR (<u>Genomics Research to Elucidate the Genetics of Rare Diseases)</u> Consortium comprising clinicians and research scientists with expertise in gene discovery, assessment of genotype-phenotype relationships, diagnostic genome analysis, variant classification, and clinical result reporting. A series of biweekly online calls were conducted over six months to draft, discuss, and refine the guidance. The group reviewed novel candidate variant/gene criteria used by several large-scale gene discovery/diagnostic efforts^38–41^ as well as descriptions from diagnostic laboratories about their evaluation and clinical reporting practices for novel candidate genes.^42–46^

### Terminology

The terms below are used as follows:

- *Clinical gene-disease validity* – an evidence-based assessment that evaluates the role of a disrupted or dysregulated gene in causing a particular Mendelian condition.^47^

Evaluating the validity of a gene’s asserted role in disease is not a binary assessment and instead, validity ranges in evidence strength. ClinGen developed a semi-quantitative framework^47^ to define evidence levels and after a delphi-survey process from the Gene Curation Coalition (GenCC)^48^, this framework was endorsed by all major gene-disease databases with harmonized assessments available in the GenCC Database (search.thegenecc.org), except for OMIM and Orphanet which use a binary inclusion or exclusion approach. While seven levels of evidence are articulated in the framework (Definitive, Strong, Moderate, Limited, No Human Disease Evidence (+/-Non-human animal model), Disputed, and Refuted), the American College of Medical Genetics (ACMG) standards indicate that the levels of Moderate to Definitive are sufficient to include in a panel clinical genetic test and that genes with Limited or less evidence should only be examined in genome-wide assessments such as ES and GS.^49^

- *Known gene* – a gene with at least a Moderate score of gene-disease validity according to the ClinGen framework (scored genes are available through GenCC^48^ (https://search.thegencc.org)
- *Novel candidate gene* – a gene in a newly-proposed gene-disease relationship. The gene either has not yet been implicated in any Mendelian condition or, more broadly, has already been implicated in a Mendelian condition but is being proposed to underlie a novel phenotype, expand the known phenotype (phenotypic expansion), or lead to a disease through a novel mechanism (e.g. different than the known mode of inheritance). These genes are a type of Gene of Uncertain Significance (GUS), which is a broader category of genes that also includes published gene-disease relationships with Limited gene-disease validity and genes that have never been implicated in disease.
- *Novel gene discovery* - the process by which variant(s) in a novel candidate gene are first identified in an affected individual, subsequently confirmed by a combination of identification of additional unrelated affected individuals with variant(s) in the same novel candidate gene and experimental studies, and finally cumulative knowledge of the new gene-disease relationship is shared with the broader community, typically via peer-reviewed publication. Additional unrelated affected individuals are most commonly identified via “matchmaking” using data sharing and networks like the Matchmaker Exchange.^50,51^
- *Candidate variant* – a variant being evaluated in the context of a novel gene-disease relationship for a causal role in a Mendelian condition. Such variants typically have a higher level of supporting evidence for functional disruption or are more rare observations (e.g. *de novo*) than variants considered in known disease genes.
- *Variant of uncertain significance* - a broad category of variation that includes variants of uncertain clinical significance found in known disease genes as well as candidate variants in novel disease genes
- *Reporting novel candidate genes* – inclusion of candidate variants in a novel candidate gene in a clinical test report provided to a patient and their clinician
- *Sharing novel candidate genes* – submission of candidate variants and phenotype data for a novel candidate gene to a repository (e.g. ClinVar) or network (e.g., Matchmaker Exchange) for purposes including facilitating future identification or “matchmaking” of additional individuals with variants in the same novel candidate gene or connection to functional investigations of the gene’s potential biological role in disease

## Results

### Criteria for novel candidate gene inclusion in clinical ES/GS

We first reviewed scoring and prioritization systems previously described by diagnostic laboratories and others^42–46,48,52^ for use in assessing the strength and types of evidence for novel candidate genes and to determine whether to share and/or report variants in the corresponding genes with the patient/provider. From this review, it was clear that evaluation of the types and strengths of evidence for novel candidate genes is just as complex, if not more so, than evaluation of evidence of variant pathogenicity. Indeed, some laboratories have developed extensive scoring and weighting systems for the evaluation of novel candidate genes,^42,45,46^ while others have more streamlined criteria or lean more heavily on statistical analysis and/or internal and external matchmaking.^43,44^

To reduce barriers to more widespread implementation of novel candidate gene identification and reporting, we focused on considering a limited set of criteria, optimized for currently computable attributes, to identify candidate genes that are most likely to become established as known disease genes in the near future. For example, 95% (150 out of 158) of new gene-disease relationships reported in 2021 and 95% (104 of 110) new gene-disease relationships reported in 2022 according to our analysis^25,53^ of discoveries described in OMIM as of May 3, 2024 satisfy at least one of these criteria (Supplementary Table 1). The 14 genes that did not satisfy any of these criteria underlie conditions that are adult-onset, have relatively mild health implications, exhibit wide variability in severity, and/or underlie an autosomal dominant condition caused only by dominant negative or biallelic variants (see footnote^a^). We recommend that variants in all novel candidate genes meeting at least one of these criteria be shared with the community by submission to the Matchmaker Exchange (MME) and ClinVar as well as considered for appropriate reporting and return to patients/providers by way of inclusion on clinical reports, as summarized in Figure 1.

**Figure 1.**
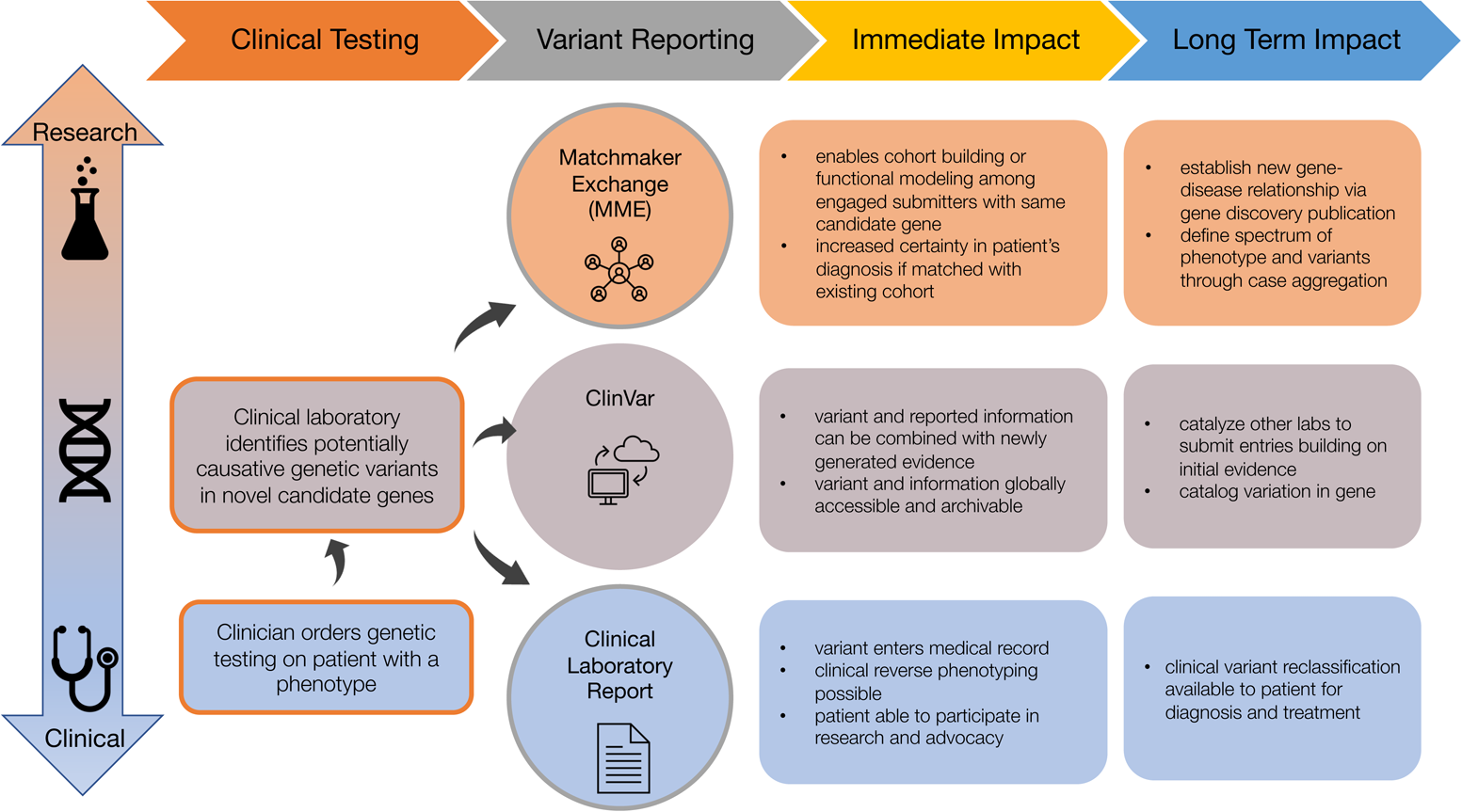
Overview of recommended venues through which laboratories can share and report novel candidate genes. Each venue (Matchmaker Exchange, ClinVar, and clinical test reports) has a different design, intended audience, and purpose, so it is not redundant to share variants in a novel candidate gene across all three. In fact, sharing across all three venues is necessary to maximize the likelihood that a novel candidate gene will be confirmed quickly.

### Recommended criteria to identify novel candidate genes to be assessed, reported, and shared

1. A gene in which predicted loss of function (pLOF) (nonsense, frameshift, essential splice site, whole or partial gene deletion or other structural variant that disrupts the coding region of the gene) or missense variant is observed to be *de novo* in a proband and the gene is strongly constrained for pLOF and/or missense variation. For proband-only analyses, strongly constrained genes with heterozygous pLOF variants absent from population databases^54^ should be considered as well. See footnote^a^ and Table 1 for further discussion of constraint.
2. A gene in which biallelic variants were identified in an individual/family with a condition seemingly inherited in an autosomal recessive pattern, and there are few or no individuals in population databases with similarly rare, predicted deleterious variants *in trans* (or phase unknown) in the gene. One resource for assessing the frequency of such individuals is the “variant co-occurrence” table in gnomAD,^55,56^ which uses in-silico phasing to predict total counts of individuals with pairs of rare missense or pLOF variants that are *in cis*, *in trans*, or with undetermined phase. Requiring at least one of the variants in the individual/family being tested to be pLOF or requiring a minimum REVEL cutoff score when the variants are missense would increase stringency of this criteria to further select only the most promising candidate genes. See Supplementary Table 1 for an example implementation.
3. A gene in which a heterozygous pLOF (including structural variant that disrupts the coding region) or missense variant was observed in a multiplex family with inheritance pattern consistent with a dominant inheritance, for which the number of segregations would reach a strong threshold of evidence,^57^ and the gene exhibits extreme constraint and is supported by functional evidence (e.g. animal model or known biological function consistent with phenotype).^52^ See footnote^a^ and Table 1 for further discussion of constraint.
4. Variant(s) in a novel candidate gene for which evidence has been gathered through collection of other cases, with or without functional studies, but the data have not yet been published. Alternatively, variant(s) in a novel candidate gene for which a convincing animal model with matching zygosity exists. A laboratory may become aware of such evidence through collection of and comparison to their internal cohort of previously tested individuals, through involvement in ongoing collaborations with researchers working on the gene, and/or other methods of scientific communication such as conference posters, talks, or preprints.
5. Variants in a known gene for which the known disease phenotype does not overlap with a proband’s clinical findings, but for which the variant-phenotype combination could represent a phenotypic expansion or novel gene-disease relationship.^58^ We recognize that such a judgement is difficult in the absence of corroborating evidence (e.g., animal model with a similar phenotype), so we recommend that such genes be reported and shared if they meet at least one of criteria 1-4.

**Table 1.**
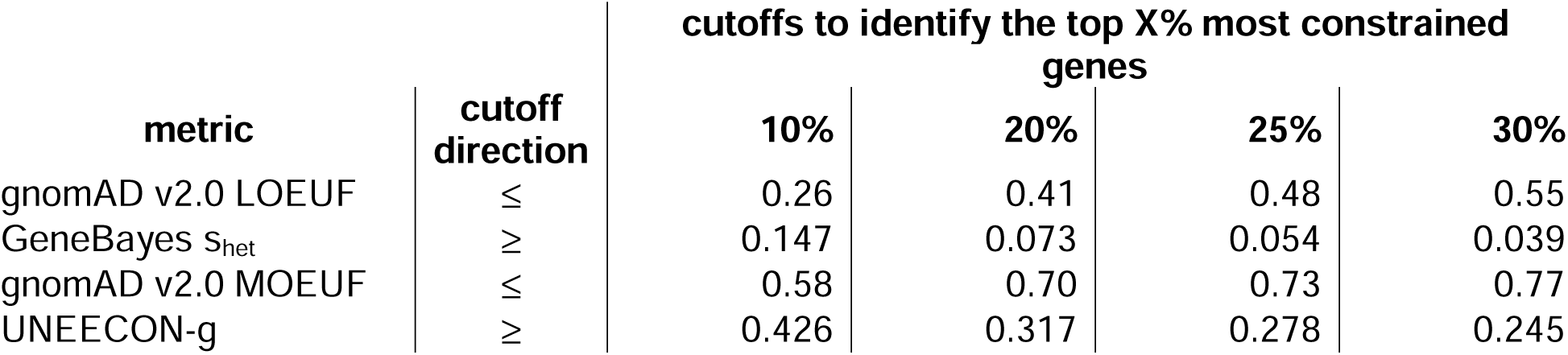
Suggested constraint metrics cutoffs for selecting novel candidate genes for criteria 1 and 3. We recommend labs apply the 20% cutoff for criteria 1 to detect strongly constrained genes and the 10% cutoff for criteria 3 to detect extremely constrained genes. However, laboratories are encouraged to consider alternative cutoffs (e.g. 25% or 30%) that would identify more candidate genes as many known genes, and therefore many novel candidate genes, are expected to fall in these ranges. In addition, note that gnomAD constraint values for LOEUF and MOEUF are not comparable across different releases (i.e. gnomAD v2.1.1 vs v4.0). We provide only gnomAD v2.0 values as the interpretation of v4.0 values is still under evaluation. See footnote^a^ for further details for interpreting constraint values.

| metric | cutoff direction | cutoffs to identify the top X% most constrained genes |  |  |  |
| --- | --- | --- | --- | --- | --- |
|  |  | 10% | 20% | 25% | 30% |
| gnomAD v2.0 LOEUF | ≤ | 0.26 | 0.41 | 0.48 | 0.55 |
| GeneBayes $s_{het}$ | ≥ | 0.147 | 0.073 | 0.054 | 0.039 |
| gnomAD v2.0 MOEUF | ≤ | 0.58 | 0.70 | 0.73 | 0.77 |
| UNEECON-g | ≥ | 0.426 | 0.317 | 0.278 | 0.245 |

### Rationale for sharing and reporting of variants in novel candidate genes

There are myriad benefits to reporting variants in novel candidate genes to clinicians and patients (and research participants) and to sharing these genes with the community despite the uncertainty of dealing with a novel candidate gene. Indeed, in terms of clinical decision-making, there may be little difference in the range of uncertainty between reporting a VUS in a known disease gene versus a candidate variant in a novel candidate gene. In both scenarios, uncertainty exists and reporting the result enables the clinician and family to collect evidence for or against a causal relationship between the variant and the patient’s phenotype. For example, reasons to routinely report VUS in known disease genes include the existence of established functional tests (e.g. metabolic assays) that can confirm or exclude the VUS as partially or fully explanatory of a patient’s clinical findings or consideration of management approaches (e.g., drug therapies, tumor surveillance, etc.) that may be low risk but of potential impact even if the VUS turns out to be benign. Similarly, routinely reporting candidate variants in novel candidate genes can encourage providers to request reanalysis from the diagnostic laboratory in the future to obtain updated information for the patient’s medical record, and in particular, if publications since the initial report have confirmed the novel candidate gene, the provider is empowered to request that the candidate variant be reclassified. It is also worth noting that both VUS^59,60^ and variants in novel candidate genes identified in underserved populations^25^ are less likely to have been previously reported let alone had evidence helpful for adjudication of pathogenicity generated. Failing to report these results risks further missed opportunities to build equity in rare disease diagnosis. Reporting a variant in a novel candidate gene also enables the provider and/or family to participate in research efforts to establish a novel gene-disease relationship through robust platforms like the Matchmaker Exchange (MME) and ClinVar that facilitate global data sharing and the ability to rapidly collect evidence in support of a candidate variant or gene via the process of “matchmaking” or “matching”.^50,61,62^

The speed and frequency with which connections to gene discovery research efforts can be made and the potential clinical impact to individual families of being connected with such research efforts should not be underestimated. Members of our group have been involved in multiple cases in which entry of the gene into MME resulted in an immediate match with a cohort described in a draft manuscript describing a novel gene discovery. Such matches enable providers and families to take advantage of results prior to publication and become engaged in research (see Box 1). These are not rare occurrences: recently, a research group described matching immediately upon submission to MME to an existing cohort with phenotypic and genetic similarities for almost ∼10% of their novel candidate genes^63^ and 15% of their submissions led to publication, with more in progress at the time of publication. Some families who received reports and were returned information about their novel candidate gene prior to publication have even launched support groups, fundraising efforts, and research into therapeutic options concurrently with researcher efforts to formally publish their novel gene discovery (e.g. *FAM177A1* [https://www.fam177a1.org]; *MAST2* and *MAST4* [https://mastgenes.org]). Thus, the benefits of identifying, reporting, and returning novel candidate variants and genes can be meaningful and immediate.

#### Box 1.

Examples in which sharing of variants in novel candidate genes enabled reporting and return of results.

A family with pontocerebellar hypoplasia and microcephaly underwent exome sequencing which identified compound heterozygosity for a missense and pLOF variant in *PPIL1*, a novel candidate gene. The variants were prioritized because they were in *trans* and one was pLOF and the gene was prioritized because no individuals in gnomAD have rare biallelic variants with at least one being pLOF. The variants and gene were submitted to MME, yielding >5 matches, including connection to investigators with a manuscript asserting a novel gene-disease relationship already under review.^64^ This quickly led to return of the novel candidate gene to the family.

A family with developmental delay and Kabuki syndrome-like facial features underwent exome sequencing which identified a *de novo* missense variant in *KDM1A*, a novel candidate gene. The *KDM1A* variant was prioritized because it was predicted pathogenic by multiple algorithms, the gene exhibits both missense and pLOF constraint, and because variants in other genes in the same pathway lead to similar phenotypes. Return of the candidate variant as a VUS to the family enabled them to create a website and engage in matchmaking themselves, followed by collaboration with researchers to publish the novel gene-disease relationship.^65,66^

In addition to benefiting individual families, efforts by laboratories to share and report novel candidate genes facilitate the process of novel gene discovery by alerting the broader community of the existence of an individual with a novel candidate gene. Such alerts are important because current gene discovery approaches are highly dependent upon matchmaking to build evidence for a candidate gene. Consequently, laboratories cannot rely upon reanalysis of past cases to trigger identification of variants in newly-published known genes, because if such genes and the variants they harbor are never shared, the gene-disease relationships may never become known.

As the number of individuals with suspected rare genetic diseases who undergo clinical testing increasingly dwarfs the number of individuals sequenced through gene discovery studies, there is a risk that the process of establishing new gene-disease relationships will slow unless novel candidate genes identified via clinical testing can be incorporated into research discovery efforts. Ultimately, facilitating gene discovery and the subsequent establishment of a new gene-disease relationship should improve diagnostic rates. For example, one clinical-research group recently reported^67^ that among 33 candidate genes/variants identified by their program, 16 were established as pathogenic and explanatory in an average of under three years, 15/33 continued to be candidates, and only 2/33 were ultimately excluded from consideration, suggesting that the yield of “real” diagnoses from novel candidate genes is high. Finally, many large-scale clinical and hybrid clinical-research genomic testing efforts are already identifying, sharing, and reporting results in novel candidate genes and have described substantial benefits in identifying a genetic diagnosis for their patients.^39,42–44,63,68^ Thus we strongly encourage clinical laboratories and research programs to proactively identify, share, and report variants in novel candidate genes.

### Guidance for sharing and reporting of variants in novel candidate genes

We encourage laboratories to adopt a tiered approach to the reporting and sharing of novel candidate genes. While we recommend that all novel candidate genes meeting at least one of the criteria described above be shared and reported, laboratories and research groups with sufficient resources^63^ and motivation may consider applying a more lenient threshold to identify and share additional variants in novel candidate genes via MME and ClinVar compared to what might be reported to an ordering provider. These genes could then be offered for return to families once further supporting evidence was available. Examples of novel candidate genes/variants that meet such a lower threshold include variants with low pathogenicity predictor scores^69^ and/or variants in genes with moderate or low pLOF/missense constraint scores^70^ in genes with known pathway/animal model evidence. Laboratories should also consider sharing genes with a recently-established gene-disease relationship as if the gene was still a novel candidate gene. Sharing knowledge that the laboratory has identified a case with a gene with only Limited or Moderate evidence for the gene-disease relationship or for which limited phenotypic information was provided in the initial gene discovery publication assists investigators seeking to build secondary cohorts to further delineate the expected phenotype and add to functional understanding.

Multiple resources support sharing of novel candidate genes (Table 2). Laboratories should at minimum share all of the following information, where supported, via each resource to facilitate efficient communication and use of the shared data for future research: gene, variant(s) (transcript ID with cDNA and amino acid change and/or genomic coordinates with genome build), zygosity (when providing case-level data), medium-to-high level structured phenotype information, and suspected mode(s) of inheritance. These data are considered minimal risk for re-identification of individuals and do not require explicit patient consent for sharing.^71–73^ Detailed rationale for the lack of direct consent requirements for sharing in ClinVar and MME are described in these referenced publications; nevertheless, these policies do encourage transparency in noting how data is shared and the importance of ongoing reanalysis and reinterpretation. Example text from clinical laboratories that can be included in test requisition forms and consent materials is included in the supplement. It is especially critical to note that when submitting to Matchmaker Exchange, the amount of effort needed to manage matching correspondence balloons enormously for all parties involved if only a gene name is shared without these additional, critical details.^63^ Similarly, one of the biggest challenges to use of ClinVar is the absence of supporting evidence and we encourage laboratories to supply the aggregate evidence directly in their submissions. Detailed guidance on what level of detail can be included in ClinVar entries without explicit consent as well as resources to support clinical laboratory data sharing can be found on the ClinGen website (https://clinicalgenome.org/share-your-data/laboratories/).

**Table 2.**
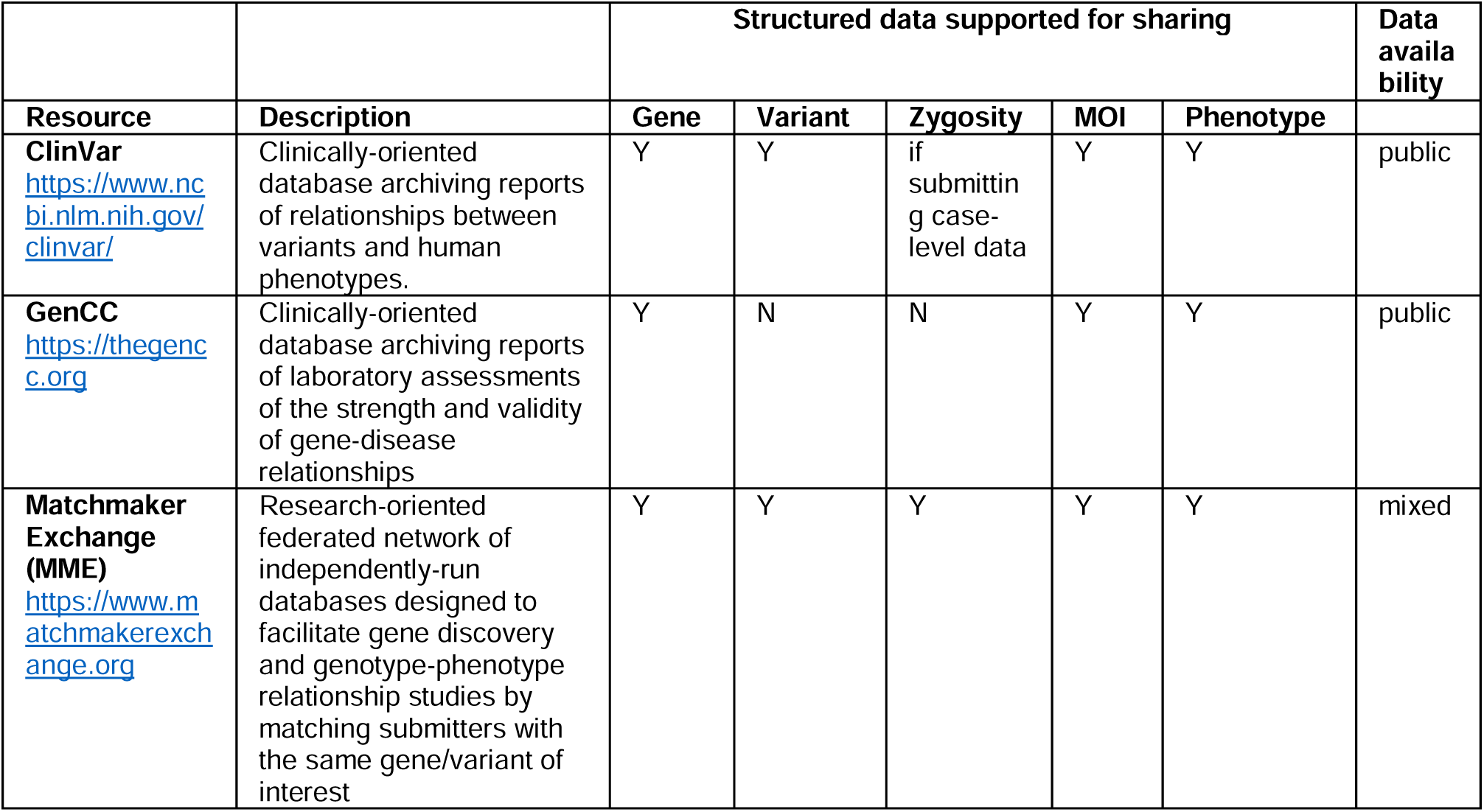
Overview of platforms that facilitate sharing of novel candidate genes. MOI = mode of inheritance.

### Guidance for choosing which platform to share candidate genes

When considering which platform to use to share candidate gene-disease relationship, several factors should be considered, as outlined here:

#### Matchmaker Exchange (MME)

The Matchmaker Exchange^50,51^ is a federated network of sites, or so-called “nodes,” run by research and clinical institutions with the goal of facilitating novel gene discovery through gene-level matching. A similar federated platform is also being developed for variant-level matching.^74^ MME nodes each support “matching” between cases based on a shared candidate gene, at minimum. Some nodes are institution-specific or have formal criteria limiting who may join to deposit case data, while three nodes (i.e. GeneMatcher, MyGene2, PhenomeCentral) are open to all self-declared clinicians and researchers. Two of these encourage (MyGene2) or allow (GeneMatcher) individuals with rare diseases or their families to join as well. We strongly encourage laboratories not yet submitting candidate genes to MME to join one of these nodes, which all have API implementations available for authorized users to facilitate integration with laboratory management systems. Use of MME is non-redundant with use of ClinVar as the users of MME are enriched for groups engaged in gene discovery and MME will alert them to the existence of each new case with a matching candidate gene.

There are two main limitations of the current MME network. First, submissions to most MME nodes are not publicly-accessible, with exceptions as described above, so other users of MME are unable to learn about interest in a candidate gene unless they have also specifically submitted that gene (i.e. if two independent submitters individually submit one of the two genes that encode binding partners in an essential protein complex, they will not be notified of the others’ submission). Second, while matching and notification of submitters is automated, follow-up to evaluate the strength and validity of matches is manual (i.e. via email) and can be time-consuming.^63^ We provide some specific recommendations to decrease the amount of communication required for engaging productively with MME (Box 2).

##### Box 2.

Recommendations for productive engagement with Matchmaker Exchange

1. Case submission:

a. Laboratories should submit their case including all of the following data elements: gene, variant(s) (transcript ID with cDNA and amino acid change and/or genomic coordinates with genome build), zygosity, and suspected mode(s) of inheritance. It is essential that medium-to-high level structured phenotype information (e.g. “intellectual disability”) is also included during submission as providing this information up front greatly reduces the number of follow-up emails that will be required to evaluate the validity of a match. Including non-identifying phenotype information in this manner does not compromise patient confidentiality^71,72^
b. Use a case-level identifier that is not identifiable but will also be recognizable to the submitter (and ideally the ordering clinician) in the future
c. Consider using a node such as DECIPHER or MyGene2 that allows submitters to make candidate genes and submission information public to increase visibility of the candidate gene-disease relationship. If submissions are not public, then the knowledge that the candidate gene has been identified will be unavailable to stakeholders not using MME to match on that exact gene (e.g. experimentalists studying the gene, geneticists with other cases due to disruptive variants in the same pathway/gene family). Other considerations may apply in choosing a node. For more guidance, see the MME website: https://www.matchmakerexchange.org/i_am_a_clinician_laboratory.html
2. Match management:

a. Submit cases to MME using a single email address/account representing the entire entity, e.g. This helps to ensure continuity as matches may not occur for years after the initial submitter may have moved on and to facilitate management of communications.
b. If responses may be delayed or conditional, the laboratory’s MME email address should have an auto-reply that details expectations about response time and the company’s policy/approach/preferences for handling inquiries about prospective matches (e.g. query submitters must provide phenotype/variant/mode of inheritance about their case up front to receive a response or query submitters must intend to participate in or collect a cohort of cases for publication, not just intend to adjudicate variants in a single case).
c. Consider notifying physicians of case submission to MME, of promising matches, and/or at minimum, note in laboratory documentation that a laboratory submits candidate genes to MME. If the node enables sharing a case with a clinician (e.g. MyGene2), doing so can circumvent having to establish a workflow for notifying clinicians.
d. Laboratories should consider modifying their test requisition forms to inform providers that if providers do not respond about promising matches within a reasonable time frame, the laboratory may reach out directly to the patient if contact info is available as facilitating establishment of their candidate gene in a novel gene-disease relationship is in the patient’s best interests.
e. If neither the patient nor clinician responds about a match that could be described in a future publication, the laboratory may consider providing a limited amount of detail (i.e. similar granularity of information that the laboratory typically submits to ClinVar) to facilitate establishment of the novel gene-disease relationship via publication.

#### ClinVar

ClinVar^62^ is a widely used public database managed by the NCBI that collects and reports variant and their phenotype associations. It is free, web accessible, and easily implemented as an attribute in variant annotation pipelines. Clinical laboratories, researchers, clinicians, and patients all use ClinVar to gather information about variants and genes, which makes it an ideal location to share variants in novel candidate genes. Its nearly ubiquitous use across clinical genomics ensures that anyone who has additional cases or information about a candidate gene would be able to make a connection or discover evidence, even if not using MME. Additionally, ClinVar features, such as providing contact information for submitters and the ability to “follow” a variant of interest further facilitating rapid data sharing and evidence building. It also has the advantage that information can more easily be exchanged without requiring additional communication through the submitter, such as occurs in MME, and therefore may be a preferred method of sharing data for low resourced laboratories.

All variants in candidate genes with limited or lower evidence should be classified and submitted to ClinVar as uncertain significance (VUS), which includes novel candidate genes, while variants in genes with moderate evidence can be classified no higher than Likely Pathogenic. To improve candidate gene variant submissions to ClinVar, we propose providing a brief summary, as follows, stating that the gene is currently a candidate for the phenotype, providing the relevant phenotypic features, noting any other relevant variants observed (e.g. those in trans) and noting the zygosity of any variant observed in the case.

##### Example language for evidence summaries when sharing a novel candidate gene in ClinVar

The {zygosity} {AminoAcidChange} variant in {Gene} was identified by our study/laboratory in [#] individual(s) with {Phenotype description}. Evidence of variant disruption includes {X}. While this gene is still lacking sufficient evidence to establish a gene-disease relationship, evidence suggests this is a possible novel gene candidate for {Disease Name/Phenotype description}. Given the limited information about this gene-disease relationship, the significance of the {AminoAcidChange} variant is uncertain. If you have any additional variant evidence or have observed other individuals with this phenotype that also have variants in {Gene} we encourage you to reach out to us.

Laboratories are encouraged to be maximally inclusive towards submitting candidate genes to MME and ClinVar to increase the chances of building more evidence supporting or refuting a candidate gene, even if the gene is not included on the corresponding clinical report.

#### Test report

Generating a clinical report describing the identification of a novel candidate gene ensures that a patient has the opportunity to benefit from the information and therefore reporting the novel candidate gene empowers them and their clinical team. The report endures as part of the medical record even if the laboratory closes, the laboratory does not proactively issue updated reports, and/or the patient’s record/data (e.g. in MME or ClinVar) are no longer maintained by the laboratory. We suggest these findings be reported in the same location of the report as other VUS but be described with language that marks them as distinct from VUS in genes with known disease relationships per the example templates below.

##### Example language for including a novel candidate gene on a report

- This variant was not identified in either of the biological parents of this individual and therefore occurred de novo. Other *de novo* variants in {Gene} have been identified in [#] individuals with {Disease Name/Phenotype description} identified through the Matchmaker Exchange (publication pending). However, this knowledgebase is still emerging and is insufficient to definitively establish the association between {Gene} and {Disease Name/Phenotype description}. In summary, additional evidence is needed to determine if {variant} may play a role in this individual’s phenotypic presentation.
- This variant was not identified in either of the biological parents of this individual and therefore occurred *de novo*. The variant is predicted to result in loss-of-function (LOF) and was identified in {Gene}, which is highly constrained for loss-of-function variants (o/e=0.1 in gnomAD), suggesting that loss-of-function variants are likely to lead to a Mendelian condition. However, no other individuals have been publicly reported with LOF variants in this gene and therefore additional evidence is needed to establish the association between {Gene} and {Disease Name/Phenotype description}. In summary, this variant occurs in a candidate gene with an uncertain role in this individual’s phenotypic presentation.

## Discussion

To date, ∼3,400 genes have at least one Moderate, Strong or Definitive gene-disease relationship in GenCC^48^ with an additional ∼1,400 listed in OMIM with at least one report of a gene-disease relationship. Additionally, twice as many (∼10,500) genes have been predicted to be novel candidate genes but currently lack a publicly asserted gene-disease relationship.^25^ However, the low prevalence of the vast majority of Mendelian conditions^75^ and increasingly widespread availability of clinical genetic testing means that many, if not most, individuals with pathogenic variants in novel candidate genes are sequenced by diagnostic laboratories alone. Together, this underscores the imperative and opportunity for diagnostic laboratories to facilitate gene discovery by assessing, sharing, and reporting novel candidate genes to clinicians, families, and researchers.

The approaches we outlined for identifying and sharing novel candidate genes, and the variants within, are conservative as we recognize that handling novel candidate genes will be new terrain for many clinical laboratories. We envision these criteria will, and should, expand over time to cover additional sets of genes and variants, as new tools and approaches for prioritizing novel candidate genes and non-coding regions and the variants within are developed. Moreover, we support laboratories with the capacity to implement more expansive criteria to do so now as we recognize that our proposed criteria prioritize stringency and ease of implementation over sensitivity to all novel candidate genes. We also encourage all groups to continue to explore emerging^50,74,76,77^ and as-yet-undeveloped approaches to efficiently share data and validate candidate genes/variants,^78^ to accelerate the process of implicating variants in new gene-disease relationships. Such engagement will increase the opportunities for even greater number of individuals with Mendelian condition to benefit from a precise genetic diagnosis.

## Acknowledgements

The GREGoR Consortium is funded by the National Human Genome Research Institute of the National Institutes of Health, through the following grants: U01HG011758, U01HG011755, U01HG011745, U01HG011762, U01HG011744, and U24HG011746. The content is solely the responsibility of the authors and does not necessarily represent the official views of the National Institutes of Health. We thank Anthony J. Marcello for assistance in data retrieval for Supplementary Table 1.

## Ethics declaration

This manuscript does not report studies on human subjects, so no IRB approval was necessary.

## Data Availability

A spreadsheet demonstrating which gene discoveries made in 2021 and 2022 would pass our criteria is provided as a supplemental table. No other original data were generated for this manuscript.

## Supplementary Materials

Supplementary Table 1 – Excel spreadsheet with multiple tabs documenting mock application of our proposed critieria to gene discoveries reported in 2021 and 2022.

Supplementary Document 1 – Word document with example text for inclusion in requisition and consent materials

Members of the Genomics Research to Elucidate the Genetics of Rare Diseases (GREGoR) Consortium:

Siwaar Abouhala^1^, Jessica Albert^2^, Miguel Almalvez^3^, Raquel Alvarez^4^, Mutaz Amin^1^, Peter Anderson^5^, Swaroop Aradhya^6^, Euan Ashley^4^, Themistocles Assimes^4^, Light Auriga^6^, Christina Austin-Tse^1^, Mike Bamshad^5^, Hayk Barseghyan^2^, Samantha Baxter^1^, Sairam Behera^7^, Shaghayegh Beheshti^7^, Gill Bejerano^4^, Seth Berger^2^, Jon Bernstein^4^, Sabrina Best^5^, Benjamin Blankenmeister^1^, Elizabeth Blue^5^, Eric Boerwinkle^8^, Emily Bonkowski^5^, Devon Bonner^4^, Philip Boone^9^, Miriam Bornhorst^2^, Harrison Brand^1^, Kati Buckingham^5^, Daniel Calame^7^, Jennefer Carter^4^, Silvia Casadei^5^, Lisa Chadwick^10^, Clarisa Chavez^4^, Ziwei Chen^4^, Ivan Chinn^7^, Jessica Chong^5^, Zeynep Coban-Akdemir^8^, Andrea J. Cohen^2^, Sarah Conner^5^, Matthew Conomos^5^, Karen Coveler^7^, Ya Allen Cui^3^, Sara Currin^10^, Robert Daber^6^, Zain Dardas^7^, Colleen Davis^5^, Moez Dawood^7^, Ivan de Dios^3^, Celine de Esch^9^, Meghan Delaney^2^, Emmanuele Delot^2^, Stephanie DiTroia^1^, Harsha Doddapaneni^7^, Haowei Du^7^, Ruizhi Duan^7^, Shannon Dugan-Perez^7^, Nhat Duong^2^, Michael Duyzend^11^, Evan Eichler^5^, Sara Emami^4^, Jamie Fraser^2^, Vincent Fusaro^3^, Miranda Galey^5^, Vijay Ganesh^1^, Brandon Garcia^7^, Kiran Garimella^1^, Richard Gibbs^7^, Casey Gifford^4^, Amy Ginsburg^5^, Page Goddard^4^, Stephanie Gogarten^5^, Nikhita Gogate^7^, William Gordon^5^, John E. Gorzynski^4^, William Greenleaf^4^, Christopher Grochowski^7^, Emily Groopman^12^, Rodrigo Guarischi Sousa^4^, Sanna Gudmundsson^1^, Ashima Gulati^2^, Stacey Hall^1^, William Harvey^5^, Megan Hawley^6^, Ben Heavner^5^, Martha Horike-Pyne^5^, Jianhong Hu^7^, Yongqing Huang^1^, James Hwang^7^, Gail Jarvik^5^, Tanner Jensen^4^, Shalini Jhangiani^7^, David Jimenez-Morales^4^, Christopher Jin^4^, Ahmed K. Saad^7^, Amanda Kahn-Kirby^3^, Jessica Kain^4^, Parneet Kaur^7^, Laura Keehan^4^, Susan Knoblach^2^, Arthur Ko^2^, Anshul Kundaje^4^, Soumya Kundu^4^, Samuel M. Lancaster^4^, Katie Larsson^1^, Arthur Lee^1^, Gabrielle Lemire^1^, Richard Lewis^7^, Wei Li^3^, Yidan Li^7^, Pengfei Liu^7^, Jonathan LoTempio^3^, James (Jim) Lupski^7^, Jialan Ma^1^, Daniel MacArthur^1^, Medhat Mahmoud^7^, Nirav Malani^6^, Brian Mangilog^1^, Dana Marafi^13^, Sofia Marmolejos^2^, Daniel Marten^1^, Eva Martinez^1^, Colby Marvin^5^, Shruti Marwaha^4^, Francesco Kumara Mastrorosa^5^, Dena Matalon^4^, Susanne May^5^, Sean McGee^5^, Lauren Meador^4^, Heather Mefford^14^, Hector Rodrigo Mendez^4^, Alexander Miller^4^, Danny E. Miller^5^, Tadahiro Mitani^15^, Stephen Montgomery^4^, Mariana Moyses^9^, Chloe Munderloh^7^, Donna Muzny^7^, Sarah Nelson^5^, Thuy-mi P. Nguyen^4^, Jonathan Nguyen^4^, Robert Nussbaum^6^, Keith Nykamp^6^, William O’Callaghan^6^, Emily O’Heir^1^, Melanie O’Leary^1^, Jeren Olsen^4^, Ikeoluwa Osei-Owusu^1^, Anne O’Donnell-Luria^1^, Evin Padhi^4^, Lynn Pais^1^, Miao Pan^2^, Piyush Panchal^7^, Karynne Patterson^5^, Sheryl Payne^5^, Davut Pehlivan^16^, Paul Petrowski^4^, Alicia Pham^1^, Georgia Pitsava^2^, Astaria/Sara Podesta^4^, Sarah Ponce^7^, Elizabeth Porter^4^, Jennifer Posey^7^, Jaime Prosser^5^, Thomas Quertermous^4^, Archana Rai^7^, Arun Ramani^6^, Heidi Rehm^1^, Chloe Reuter^4^, Jason Reuter^6^, Matthew Richardson^5^, Andres Rivera-Munoz^7^, Oriane Rubio^4^, Aniko Sabo^7^, Monica Salani^1^, Kaitlin Samocha^1^, Alba Sanchis-Juan^1^, Sarah Savage^6^, Evette Scott^7^, Stuart Scott^4^, Fritz Sedlazeck^7^, Gulalai Shah^1^, Ali Shojaie^5^, Mugdha Singh^1^, Kevin Smith^4^, Josh Smith^5^, Hana Snow^1^, Michael Snyder^4^, Kayla Socarras^1^, Lea Starita^5^, Brigitte Stark^4^, Sarah Stenton^12^, Andrew Stergachis^5^, Adrienne Stilp^5^, V. Reid Sutton^16^, Jui-Cheng Tai^9^, Michael (Mike) Talkowski^1^, Christina Tise^4^, Catherine (Cat) Tong^5^, Philip Tsao^4^, Rachel Ungar^4^, Grace VanNoy^1^, Eric Vilain^3^, Isabella Voutos^4^, Kim Walker^7^, Chia-Lin Wei^5^, Ben Weisburd^1^, Jeff Weiss^5^, Chris Wellington^10^, Ziming Weng^4^, Emily Westheimer^6^, Marsha Wheeler^5^, Matthew Wheeler^4^, Laurens Wiel^4^, Michael Wilson^1^, Monica Wojcik^1^, Quenna Wong^5^, Changrui Xiao^3^, Rachita Yadav^1^, Qian Yi^5^, Bo Yuan^7^, Jianhua Zhao^6^, Jimmy Zhen^4^, Harry Zhou^6^

^1^Broad Institute, ^2^Children’s National, ^3^University of California Irvine, ^4^Stanford University, ^5^University of Washington, ^6^Invitae, ^7^Baylor College of Medicine, ^8^University of Texas Health Science Center, 9Massachusetts General Hospital; Broad Institute, ^10^National Institutes of Health (NIH), ^11^Broad Institute; Massachusetts General Hospital; Boston Children’s Hospital, ^12^Broad Institute; Boston Children’s Hospital, ^13^Kuwait University, ^14^St. Jude Children’s Research Hospital, ^15^Jichi Medical University, ^16^Baylor College of Medicine; Texas Children’s Hospital

## Footnotes

a Gene constraint is a measure of the degree of deficit of a type of variant (e.g. pLOF, missense) among individuals in a population database, where a greater deficit is suggestive of a gene with little tolerance for the variant type. In gnomAD, the observed/expected ratios^54^ for pLOF (LOEUF), missense (MOEUF), and synonymous variants are provided along with other measures of constraint (pLI,^79^ Z scores). GeneBayes s_het_ (bioRxiv doi:10.1101/2023.05.19.541520) and UNEECON-g^80^ are two other estimates of gene-level strength of selection against LOF variants and missense variants, respectively, that use machine learning to enable more sensitive predictions of constraint.^80^ Fine-grained measures of constraint can detect subgenic/regional [Gnocchi^81^, Constrained Coding Regions (CCR)^82^] or even single amino acid [MPC (bioRxiv doi:10.1101/148353), HMC (medRxiv doi:10.1101/2022.02.16.22271023)] constraint. Constraint metrics are most sensitive for genes when the pathogenic mechanism is LOF, the condition is early-onset, and the condition has at least moderate effects on reproductive fitness. In contrast, these metrics have reduced sensitivity when the pathogenic mechanism is typically dominant negative (DN) or gain-of-function (GOF) effects,^83,84^ which are often due to missense variants but can also be caused by variants that appear to be pLOF but escape nonsense-mediated decay.^85,86^ Such genes (e.g. *HRAS, TRPC6, POLR3B*, *PRSS1*, *HSPB1*) often do not exhibit constraint for pLOF or missense variation according to some or even all currently-available metrics as listed in criteria 1, and GOF and DN variants are also often not predicted as damaging by computational variant effect predictors. pLOF variants in novel candidate genes should be examined carefully as such variants are more likely to not result in true LOF.^87^ For example, the variant may be found in an exon that is alternatively transcribed and not required for function.^54,88^ Genes that underlie conditions with modest or low effect on reproductive fitness (e.g. *C9ORF72* underlying amyotrophic lateral sclerosis and *TYR* underlying oculocutaneous albinism are not constrained for pLOF or missense variation) also exhibit less constraint.

