## Supplemental Document 1 for "Considerations for reporting variants in novel candidate genes identified during clinical genomic testing"

**Supplementary Document**

Detailed rationale for the lack of direct consent requirements for sharing [candidate] gene, variant(s) (transcript ID with cDNA and amino acid change and/or genomic coordinates with genome build), zygosity (when providing case-level data), medium-to-high level structured phenotype information, and suspected mode(s) of inheritance in ClinVar and MME have been described previously.^1–3^ Nevertheless laboratories are encouraged to be transparent about how data is shared and the importance of ongoing reanalysis and reinterpretation. Example text from clinical laboratories that can be included in test requisition forms and consent materials with this goal in mind follows:

**GeneDx**

DATABASE PARTICIPATION De-identified health history and genetic information can help healthcare providers and scientists understand how genes affect human health. Sharing this deidentified information helps healthcare providers to provide better care for their patients and researchers to make new discoveries. GeneDx shares this type of information with healthcare providers, scientists, and healthcare databases. GeneDx will not share any personally identifying information and will replace the identifying information with a unique code not derived from any personally identifying information. Even with a unique code, there is a risk that I could be identified based on the genetic and health information that is shared. GeneDx believes that this is unlikely, though the risk is greater if I have already shared my genetic or health information with public resources, such as genealogy websites.

**Ambry**

Matchmaker Exchange: Ambry participates in the Matchmaker Exchange through GeneMatcher to facilitate novel disease-gene discovery through connecting health care providers with other external clinicians and researchers. This process has led to over 100 collaborations and over 40 publications.

**Mass General Brigham Laboratory for Molecular Medicine**

*Will anyone else have access to my genomic sequence, shared medical history or interpreted results?* The ordering physician can obtain access to your genomic sequence data files for the purpose of your clinical care. Test results and submitted clinical information may be shared with other clinical laboratories for the purpose of improving our understanding of the relationship between genetic changes and clinical symptoms. Sharing data in this manner may enable us to provide better interpretations of your genetic findings as well as assist other patients with similar results. We will protect your privacy/confidentiality by replacing your name and other direct identifiers, such as date of birth or medical record number, with a code. The key to the code numbers will be stored securely in the testing laboratory. We will share only de-identified information with outside clinical labs.

1. Azzariti DR, Riggs ER, Niehaus A, et al. Points to consider for sharing variant-level information from clinical genetic testing with ClinVar. *Mol Case Stud*. 2018;4(1):a002345. doi:10.1101/mcs.a002345

2. Dyke SOM, Knoppers BM, Hamosh A, et al. “Matching” consent to purpose: The example of the Matchmaker Exchange. *Hum Mutat*. 2017;38(10):1281-1285. doi:10.1002/humu.23278

3. Wright CF, Ware JS, Lucassen AM, et al. Genomic variant sharing: a position statement. *Wellcome Open Res*. 2019;4:22. doi:10.12688/wellcomeopenres.15090.2
